# Absence of Telomerase Leads to Immune Response and Tumor Regression in Zebrafish Melanoma

**DOI:** 10.1101/2023.03.24.534079

**Authors:** Bruno Lopes-Bastos, Joana Nabais, Tânia Ferreira, Mounir El Maï, Malia Bird, Seniye Targen, Da Kang, Jia-Xing Yue, Tânia Carvalho, Miguel Godinho Ferreira

## Abstract

Most cancers reactivate telomerase to maintain telomere length to acquire immortality. The importance of this process is well illustrated by the fact that telomerase promoter mutations are found at a high frequency in many cancer types, including melanoma. However, it is unclear when and if telomerase is strictly required during tumorigenesis. Here, we show that melanoma can occur in the absence of telomerase but is required to sustain later growth and to avoid tumor regression. We combined telomerase mutant zebrafish (*tert-/-*) with two established melanoma models and found equal melanoma incidence and invasiveness as tumors became visible. Later, however, while *tert+/+* fish develop increasing larger tumors, *tert-/-* tumors stagnate growth and regress. *tert-/-* tumors showed lower cell proliferation, higher apoptosis and melanocyte differentiation. We also detected an immune response directed at *tert-/-* tumors. *tert-/-* tumors exhibited increased immune cell infiltrates and resume growth when transplanted into immunocompromised hosts. We propose that telomerase is required for melanoma in zebrafish, albeit at later stages of progression, to sustain growth while avoiding immune rejection and regression. Thus, absence of telomerase restricts melanoma through tumor-autonomous mechanisms (cell cycle arrest, apoptosis and melanocyte differentiation) and a non-tumor-autonomous mechanisms (immune rejection).

## Introduction

In most adult somatic tissues, telomeres shorten through successive cell divisions. Short telomeres cause telomere dysfunction, leading to replicative senescence and cell cycle arrest^1–4^. Telomere shortening imposes two proliferation barriers that pre-cancerous cells must overcome to become fully malignant^5,6^. The first proliferation barrier, known as M1 or senescence, is composed of a p53-dependent mechanism, overcome by loss of p53/Rb function. The second, known as M2 or crisis, is characterized by increased genome instability fueled by telomere-to-telomere fusions^5^. To reverse this state, cells must engage in a telomere maintenance mechanism (TMM). Approximately 90% of human tumors re-activate telomerase, thus elongating telomeres and chromosome-end protection^7^. Telomerase is composed of two core components, a reverse transcriptase DNA polymerase subunit (TERT) and a template containing noncoding RNA subunit (TR). Telomerase binds to the 3’ end of telomeres and processively adds (TTAGGG)n hexanucleotide repeats^8^. In the absence of telomerase, cancer cells can maintain telomeres through a recombination-based mechanism known as alternative lengthening of telomeres (ALT)^6,9^. *In vivo* studies using murine models showed that the lack of telomerase, in animals with short telomeres, results in less cancer formation^10,11^. Additionally, telomerase inhibition of established tumors resulted in growth restriction^10^. Interestingly, tumor relapse upon telomerase inhibition was observed in some animals in which ALT was detected^10^, showing that ALT can be an efficient TMM in tumorigenesis. ALT was also observed in zebrafish brain tumors^12^.

In contrast to the previous idea, a study that analyzed 31 cancer types, failed to detect TMMs in 22% of 6,835 human samples^13^. In addition, tissue culture of cells derived from melanoma and neuroblastoma TERT-/ALT-tumors displayed progressive telomere shortening, confirming those tumors lacked an active TMM in patients^14,15^. Cancers lacking TMMs come at a cost that limits their replicative capacity imposed by the Hayflick limit. This trade-off is illustrated by neuroblastoma lacking TMMs. These cancers present a better prognosis and are correlated with spontaneous regression^16^. However, this is not always the case. Melanoma lacking a TMM were reported to be extremely aggressive, despite limited proliferative capacity in cell culture^15^. In addition, TMMs may play a role in resistance to immunotherapies. Studies have shown that metastatic melanoma resistant to immune checkpoint blockade therapies displays an increased expression of genes related to telomere maintenance^17^, while TERT mutations can predict the response to some immune checkpoint blockade therapies^18,19^. These findings suggest that telomerase expression may regulate cancer immunogenicity, potentially serving as a predictive biomarker for therapy.

Replicative immortality provided by TMMs is a well-established hallmark of cancer. However, it’s unclear when during tumorigenesis TMMs become activated and whether they are strictly essential for lethal cancers. Activating TERT promoter mutations (TPM) are found at a high frequency in several of human tumor types^20–23^. These tumors include cutaneous melanomas at a frequency of 60-85%^20,21,24^. TPMs have been identified in both early- and late-stage tumors^20–23,25^, suggesting that telomerase activation may occur throughout tumorigenesis. Here, we investigated the role of telomerase in melanoma initiation and progression with the goal of identifying at which stage TMMs are strictly required for tumorigenesis. Unexpectedly, we observed that melanoma initiation and early progression does not require telomerase nor ALT. However, lack of telomerase activity at later stages restricted melanoma growth and led to tumor regression through both tumor-autonomous and non-tumor-autonomous mechanisms.

## Results

### Melanoma initiation and early progression does not require TMMs

To determine the role of telomerase in melanoma, we crossed the telomerase mutant zebrafish (tert^hu3430^, hereby referred as *tert-/-*), in which telomerase is enzymatically inactive^26^, with the *Tg(mitfa:Hsa.HRAS_G12V,mitfa:GFP)* melanoma model, referred here as mitfa:HRAS, developed by the Hurlstone lab^27^. Melanocytes of mitfa:HRAS fish were shown to exhibit hyperplasia, dysplasia and to spontaneously progression to invasive melanoma^27^. In agreement, we observe that wildtype telomerase (*tert+/+*) developed melanoma, reaching full penetrance by 4 months of age (Figure 1A). In this model, the transgene also induces the expression of GFP under the *mitfa* promoter to identify tumor development (Figure 1B). Surprisingly, lack of telomerase activity in telomerase mutant fish harboring mitfa:HRAS (mitfa:HRAS; *tert-/-*) did not affect the dynamics of melanoma incidence (Figure 1A). Thus, telomerase is not required for melanoma formation in this model. This result differs from previous data indicating that late-generation telomerase-deficient mice (G5 terc-/-) are resistant to skin carcinoma^11^.

**Figure 1:**
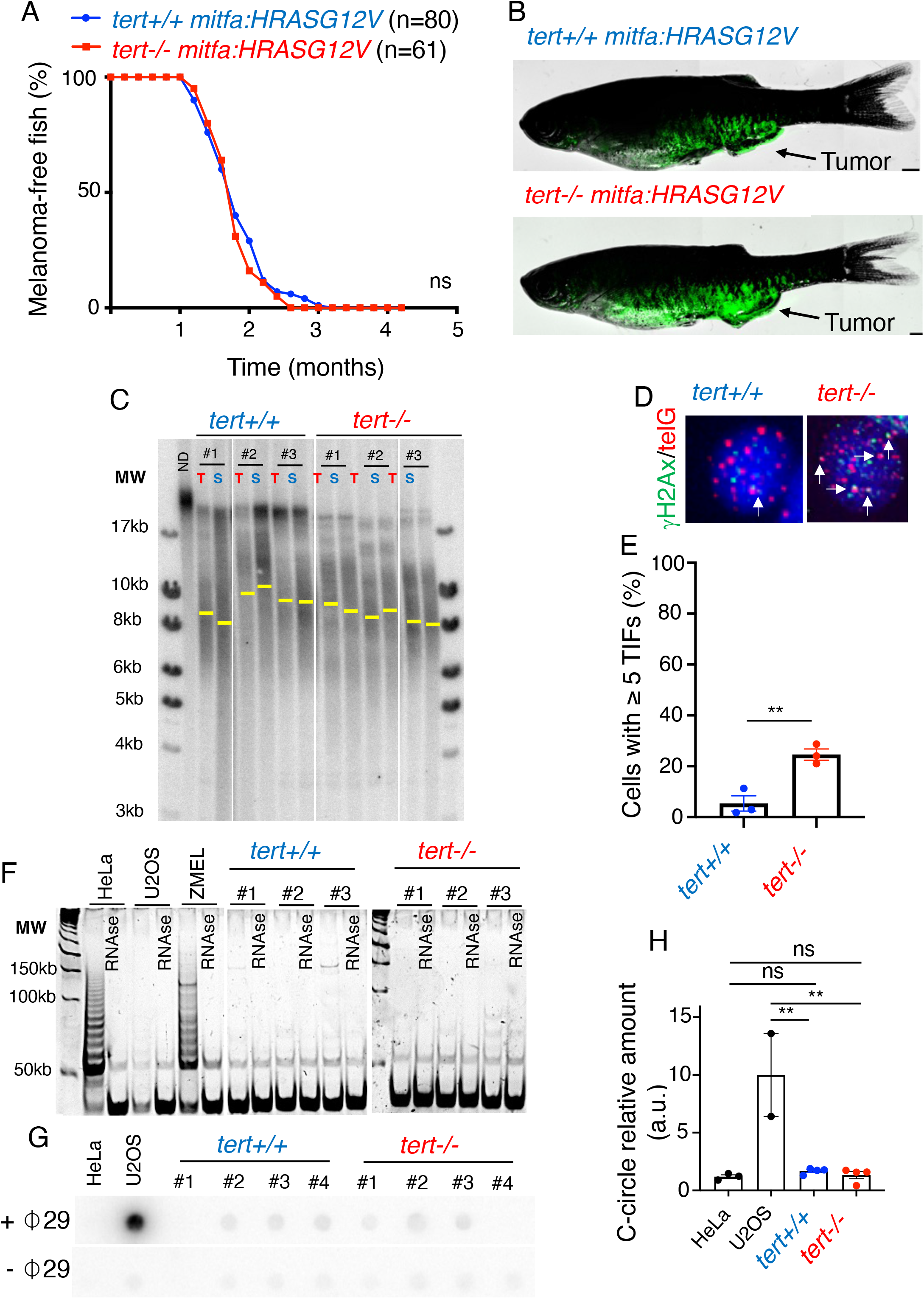
Melanoma initiation does not require a telomere maintenance mechanism. (A-B) Incidence of melanoma of mitfa:HRAS fish is not affected by the lack of telomerase. A-Percentage of tumor-free fish compared between *tert+/+* and *tert-/-* fish (log-rank test). B-Representative images of *tert+/+* and *tert-/-* fish with melanoma expressing GFP. (C-D) Telomere shortening is not apparent between *tert+/+* and *tert-/-* fish. However, *tert-/-* tumors possess higher DNA damage at telomeres. C-Telomere restriction fragment (TRF) analysis by Southern blotting of tumor (T) and skin (S) genomic DNA extracted from 3-month-old fish and quantifications for mean telomere length (yellow bars). D-Images of γH2Ax/telG immune-FISH of tumor cells derived from *tert+/+* and *tert-/-* 3-month-old fish. White arrows indicate Telomere Dysfunction-Induction Foci (TIF) in tumor cells. E-Percentage of cells containing ≥5 TIFs from D (n=3; unpaired *t*-test). (F-H) Early-stage melanoma do not display TMMs. F-Telomerase activity evaluated by TRAP of tumor samples derived from *tert+/+* and *tert-/-* 3-month-old fish (n=3). Extracts from HeLa and ZMEL cells were used as positive controls and U2OS as a negative control. G-C-circle assay of tumor samples derived from *tert+/+* and *tert-/-* 3-month-old fish. Extracts from HeLa cells and U2OS were used as negative and positive controls, respectively. H-Quantification of C-circle signal (n=4; One-way ANOVA). Error bars represent ±S.E.M.; each dot represents an individual tumor; \*\**p*≤0.01; ns= not significant.

To extend our findings, we combined the telomerase mutant fish with a second widely used melanoma model expressing the human *BRAFV600E* oncogene in a *tp53* deficient background, *Tg(mitfa:BRAF^V600E^); tp53*^*-/-*28^. Similarly, we observed that *Tg(mitfa:BRAF^V600E^); tp53^-/-^; tert-/-* does not affect melanoma incidence (supplementary Figure 1A). This second melanoma model also allowed us to observe the different phases of melanogenesis. In both *tert+/+* and *tert-/-* we observed the three different stages of melanoma progression: benign lesion, radial growth phase and vertical growth phase, (supplementary Figure 1B). Our results show that, even in the absence of telomerase, melanoma progresses efficiently through all these stages. Even though BRAFV600E is the most frequent alteration in melanoma, we chose to conduct the remaining of our experiments using the mitfa:HRAS melanoma model. In contrast to the BRAFV600E model, the mitfa:HRAS zebrafish do not require a pre-existing *tp53* mutation to develop tumors^27^. We and others have shown that p53 dysfunction suppresses *tert-/-* and *tert-/-* p53-/- double mutants do not display the phenotypes associated with lack of telomerase^26,29^. Therefore, the BRAFV600E model would not allow us to directly investigate the function of telomerase in tumorigenesis. It is worth noting that Ras mutations, even though less frequent, are still present in human melanoma and in need of effective therapies^24^.

In order to evaluate whether telomerase has an impact in melanoma initiation, we studied the incidence of early tumorigenesis in mitfa:HRAS; *tert-/-* fish. Our results show that lack of telomerase did not affect either the development of microtumors in 1-2-month-old fish (supplementary Figure 1C-D) or their invasiveness capacity (supplementary Figure 1E-F). The similar incidence of early tumors in telomerase mutant fish reinforces the idea that telomerase activation is not required for cancer initiation.

Next, we analyzed visible tumors of 3 months old fish. We defined this age as an early progression stage in tumorigenesis given that almost all fish carry macroscopic lesions (Figure 1A). We measured telomere length using Telomere Restriction Fragment (TRF) Southern blotting and compared telomeres from tumor samples to those derived from the skin of the same fish, as internal controls. Consistent with our previous results, telomeres of *tert-/-* fish are shorter that *tert+/+* (Figure 1C). Unexpectedly, we failed to detect telomere shortening in tumors compared to skin from either *tert+/+* or *tert-/-* 3 months old fish (Figure 1C-D). Nevertheless, *tert-/-* melanoma cells exhibited a significant increase in telomere damage induced foci (TIFs) compared to *tert+/+* tumor cells (*tert+/+*: 5% versus *tert-/-*: 25%; N=3 *p*=0.0068, Figure 1D-E). Thus, even though *tert-/-* melanomas show similar average telomere length compared to skin, our result show that *tert-/-* melanoma cells harbor critically short telomeres even at this early progression stage.

Most cancers present activating telomerase promoter mutations from early stages of carcinogenesis^30^. To evaluate whether tumors in our model present telomerase activity, we performed a Telomerase Repeated Amplification Protocol (TRAP) assay. We used human HeLa cells and zebrafish melanoma ZMEL cell line as a positive controls^31^. As negative control, we used the osteosarcoma U2OS cell line. As expected, zebrafish *tert-/-* tumors did not show telomerase activity (Figure 1F). Surprisingly, early *tert+/+* tumors also failed to show detectable telomerase activity. This result suggests that zebrafish melanoma does not activate telomerase at an early stage of tumorigenesis.

Cancer cells can also maintain telomeres by engaging in an alternative mechanism to telomerase known as Alternative Lengthening of Telomeres (ALT). ALT cells are characterized by an increase in the heterogeneity of telomere length, displaying longer and shorter telomeres than TERT-positive cells^32^. However, analyzing TRFs derived from both *tert+/+* and *tert-/-* tumors, we observed a similar spread of telomere sizes (Figure 1C). ALT is also characterized by the presence of extrachromosomal DNA, known as C-circles^33^. Amplification of telomeric circles using Φ29 DNA polymerase allows for amplification of C-circles and provides a robust way to assess ALT-positive cells, such as U2OS^33^. However, this assay failed to detect the presence of C-circles in *tert-/-* tumors (Figure 1G-H). Consistently, a previous study using Ras-induced zebrafish melanoma also did not exhibit an increase in C-circles^12^. Altogether, our data shows that melanoma initiation and early progression does not require telomerase and that these tumors do not engage in ALT to maintain telomere length.

### Lack of telomerase leads to growth restriction in late progression

Our observation that telomerase activation was not required for melanoma early progression prompted us to investigate its effect on later stages. We followed tumor progression over time and assessed tumor area in fish aged 5, 7 and 9-months-old (Figure 2A-B; Supplementary Figure 2A). In 5-month-old fish, the average tumor area was indistinguishable between *tert+/+* and *tert-/-* fish. However, while *tert+/+* tumor size increased overtime, the average area of *tert-/-* tumors remained stable and was significantly smaller than *tert+/+* tumors (*p*= 0,0269 for 7 months; *p*= 0,0017 for 9 months). Tumor size differences were more evident when individual tumor trajectories were taken into consideration (Figure 2C). All *tert+/+* tumors increased in size overtime, while the size of most *tert-/-* tumors remained constant (Figure 2C). By calculating the regression slope for each tumor size variation (Figure 2D), we found that all *tert+/+* tumors had a positive trend while most *tert-/-* tumors either did not grow (slope ≈ 0) or regressed. This result indicates that *tert-/-* tumors exhibited slower growth dynamics at later stages.

**Figure 2:**
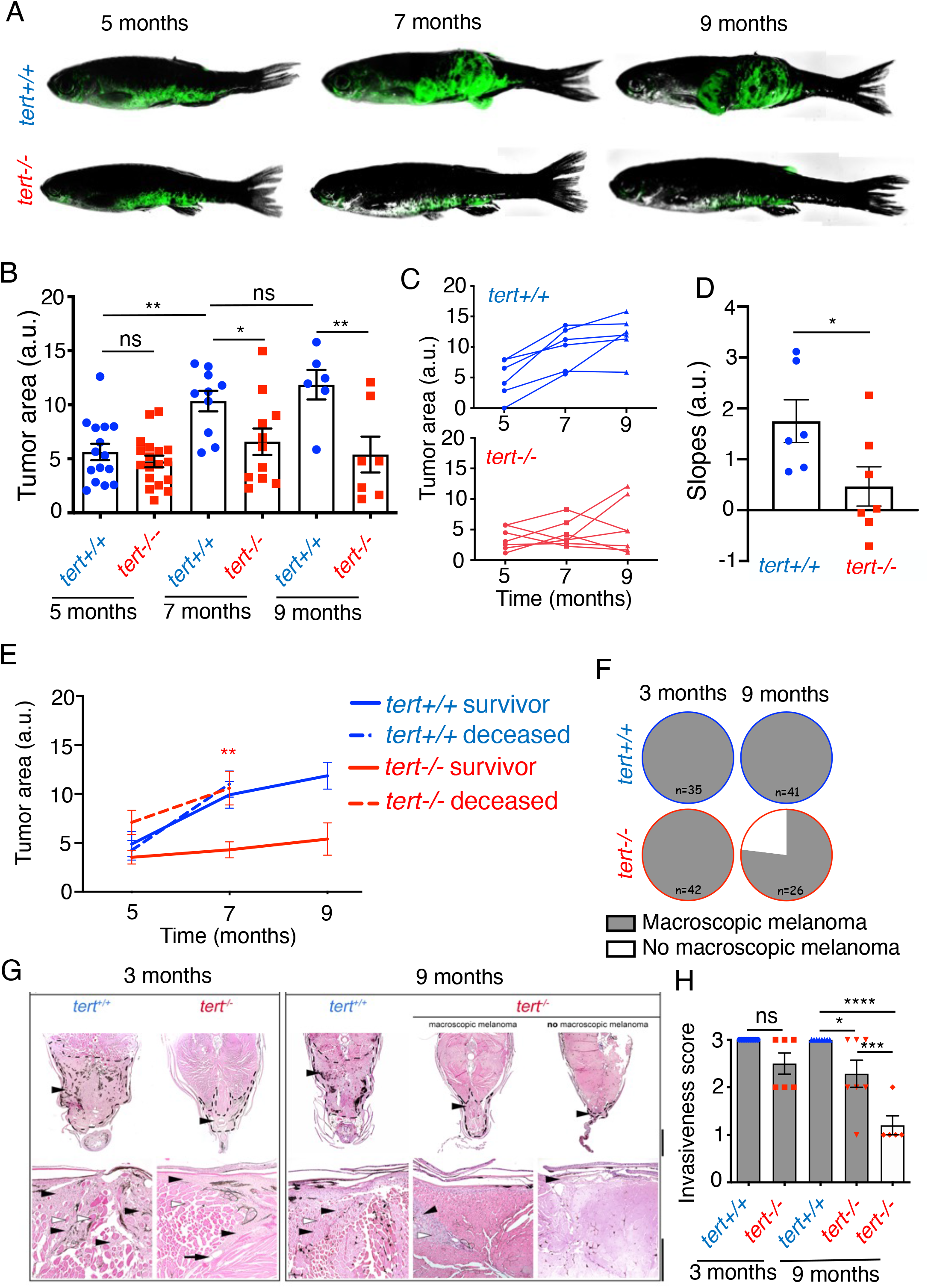
Absence of telomerase restricts tumor growth leading to melanoma regression. (A-D) Lack of telomerase impacts late tumor growth. A-Examples of melanoma evolution overtime of *tert+/+* and *tert-/-* fish. B-Quantification of tumor size of 5-, 7- and 9-month-old *tert+/+* and *tert-/-* fish (One-Way ANOVA). C-Evolution tumor size overtime in individual animals (n=6 and n=7). D-Slope of tumor size evolution calculated using linear regression of three time-points (unpaired *t*-test). E-Larger tumors are associated with *tert-/-* fish lethality. Tumor size evolution of fish that either survived until 9 months (solid line: survival group) or died after 7 months (dashed line: non-survivor group). Statistics compare *tert-/-* survivors with *tert-/-* deceased fish at 7 months (unpaired *t*-test). (F-H) Melanoma regresses in fish lacking telomerase. F-*tert+/+* and *tert-/-* fish with macroscopic tumors at 3 and 9 months of age. G-Histopathology analysis of melanoma of 3- and 9-month-old *tert+/+* and *tert-/-* zebrafish. Tumors are indicated by a black arrowhead and dashed line. Higher magnification shows infiltrative features of melanoma (black arrowhead), with marked invasion, destruction, and replacement of the hypaxialis muscle (white arrowhead). Top scale: 500 μm and bottom scale: 200 μm. H-Quantification of melanoma invasiveness by 1-non-invasive, 2-minimally invasive, 3-invasive. Error bars represent ±S.E.M.; each dot represents an individual tumor; * *p*≤0,05; ** *p*≤0,01; *** *p*≤0,001, \*\*\*\**p*≤0,0001; ns: not significant; a.u. arbitrary units.

Although *tert-/-* tumors were smaller in size, they were as lethal as *tert+/+* tumors (Supplementary Figure 2B). We previously reported that *tert-/-* fish age prematurely and exhibit reduced survival^26^. This effect could explain the high mortality of the *tert-/-* fish rather than the melanoma per se. To examine this reduced survival further, we analyzed the survival of cousins of the fish used in this study as a control (Supplementary Figure 2B-C). Survival of fish lacking mitfa:HRAS was close to 100% throughout the experiment and *tert-/-* did not differ from *tert+/+* fish. This suggests that the high mortality observed in mitfa:HRAS *tert-/-* fish is not due to premature aging but rather a consequence of the melanoma itself. Given the differences in tumor size, we also tested whether size could predict melanoma survival rates. To answer this question, we quantified the animals that survived until 9 months of age (“survivor” group) and the ones that died between the 7 and 9 months (“deceased” group) for each given genotype (Figure 2E). We observed that tumor size did not differ between the survivor and the deceased groups in *tert+/+* fish, indicating that tumor size did not predict survival for *tert+/+* animals (dashed and filled lines of *tert+/+* fish in Figure 2E). In contrast, the non-survival group of *tert-/-* fish possessed larger tumors than the survivor group (*p*= 0,0044, dashed and filled lines of *tert-/-* fish in Figure 2E). This result indicates that tumor size greatly impacts survival of *tert-/-* fish. It also shows that *tert-/-* tumors are capable of growing to the same extent as those of *tert+/+* fish (Figure 2E). However, in contrast to *tert+/+, tert-/-* fish carrying larger tumors die and are excluded from later stages of melanoma progression.

Our previous results showed that, by 3-months of age, nearly all *tert+/+* and *tert-/-* fish possessed macroscopic tumors (Figure 2F) that were histologically comparable with similar invasiveness profiles (Figure 2G-H). In contrast, by 9-months of age, while all *tert+/+* tumors still exhibited clear tumors, a significant proportion of *tert-/-* fish (ca. 25%, N=26) lacked visible tumors, suggestive of melanoma regression (Figure 2F). We were still able to find histological remains of melanoma with reduced invasiveness in these fish (Figure 2G-H). Muscle fibers of regressing melanoma fish displayed signs of atrophy with empty spaces between the fibers. A possible explanation could be that melanoma invaded the muscle and when it regressed, the muscle fibers did not recover. The remaining macroscopic *tert-/-* tumors displayed significantly reduced invasiveness when compared to *tert+/+* tumors (*p*= 0,0257, Figure 2G-H). Overall, our results suggest that *tert-/-* tumors can grow as much as *tert+/+* tumors, but they result in earlier mortality relative to *tert+/+* fish. Thus, telomerase not only is required to sustain tumor grow in late melanoma progression, but lack of telomerase increases the susceptibility of the organism to larger tumors.

### Telomerase is re-activated in late melanoma progression

Given that late melanoma progression is defective upon telomerase deficiency, we sought to determine whether telomerase is re-activated in tumors of *tert+/+* fish. We assayed telomerase activity by TRAP in 9 months old fish and, indeed, we were able to detect telomerase activity in *tert+/+* tumors but not, as expected, in *tert-/-* tumors (Figure 3A). Thus, telomerase is re-activated in melanoma cells but only after tumors are formed and macroscopically visible. We next analyzed telomere length in tumors derived from 7 months old fish using by TRF (Figure 3B). We observed that, while telomeres of *tert-/-* tumors are shorter than their matched skin controls, *tert+/+* tumors maintained an average telomere length similar to the skin (Figure 3B). Consistently, *tert-/-* tumors displayed greater levels of DNA damage foci at telomeres (*p*= 0,0421, Figure 3C-D) than those observed in early-stage tumors (Figure 1D-E). It is striking that, even in the absence of telomerase, *tert-/-* fish possessed tumors by 9 months of age. In order to verify whether these tumors developed ALT as TMM, we looked for the presence of C-circles. Surprisingly, we did not obtain Φ29 DNA polymerase amplification of C-circles in neither *tert+/+* nor *tert-/-* tumors (Figure 3E-F), denoting that mitfa:HRAS tumors do not activate ALT at later progression stages. Altogether, as expected from other models, telomerase is indeed expressed during melanoma progression in telomerase proficient individuals. However, we highlight that telomerase activation is a late event in tumorigenesis and that it occurs upon telomere shortening and increased DNA damage at telomeres.

**Figure 3:**
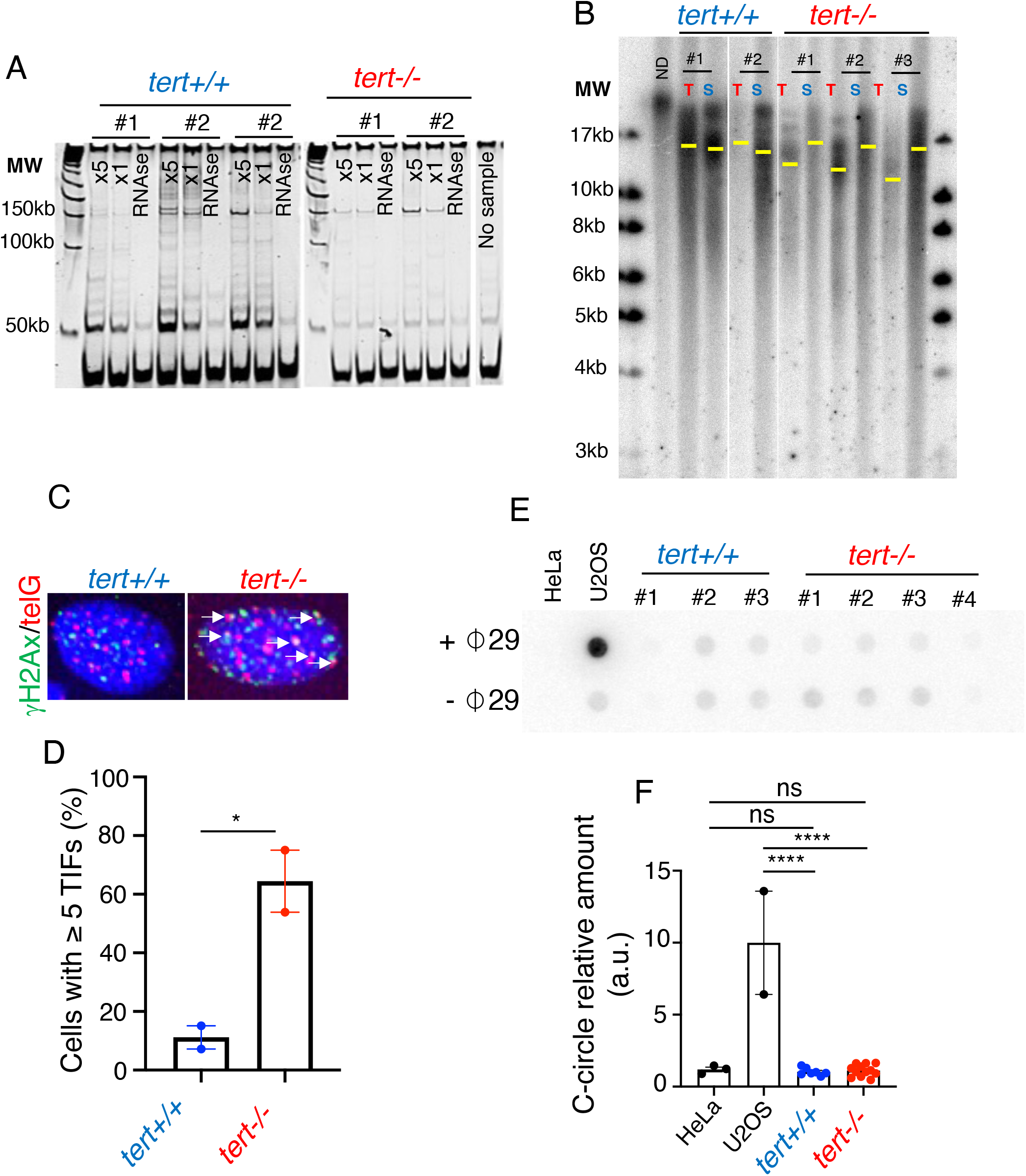
Telomerase is re-activated in late-stage melanoma and its absence causes telomere shortening. (A-B) Telomerase is active in tumors of 9-month-old fish and telomere shortening occurs in *tert-/-* tumors. A-Telomerase activity evaluated by TRAP in tumor samples derived from *tert+/+* and *tert-/-* 9-months old fish (n=3 and n=2, respectively). B-Telomere restriction fragment (TRF) analysis by Southern blotting of tumor (T) and skin (S) genomic DNA extracted from 7-month-old fish and quantifications for mean telomere length (yellow bars). C-Images of γH2Ax/telG immune-FISH of tumor cells derived from *tert+/+* and *tert-/-* 9-month-old fish. White arrows indicate Telomere Dysfunction-Induction Foci (TIF) in tumor cells. D-Percentage of cells containing ≥5 TIFs from B (n=2; unpaired *t*-test). (E-F) Late melanoma in *tert-/-* fish do not engage ALT as TMM. E-C-circle assay of tumor samples derived from *tert+/+* and *tert-/-* 3-month-old fish. Extracts from HeLa cells and U2OS were used as negative and positive controls, respectively. F-Quantification of C-circle signal (n≥7, One-way ANOVA). Error bars represent ±S.E.M.; each dot represents an individual tumor; * *p*≤0,05; \*\*\*\**p*≤0,0001; ns: not significant.

### Tumor-autonomous causes for *tert-/-* tumor growth decline

We showed that lack of telomerase leads to telomere shortening and growth restriction in late melanoma progression. To understand the mechanism behind the restrained growth of *tert-/-* tumors, we compared the expression profiles of *tert+/+* vs *tert-/-* tumors using RNAseq using early stage (3-months-old) and later stage (9-months-old) melanoma tumor samples. GSEA analysis showed that proliferation pathways are down-regulated in *tert-/-* tumors in 9-month-old fish while apoptosis is up-regulated (Figure 4A). We sought to confirm these results using immunofluorescence. We observed that DNA damage is increased in *tert-/-* melanoma at later stages (γH2Ax, *p*= 0,0054, Figure 4B). Consistently, we observed that cell proliferation rates are lower (PCNA, *p*<0,0001, Figure 4C) while apoptosis levels are higher (cleaved-caspase-3, *p*=0,005, Figure 4D) in later stage *tert-/-* melanoma. In contrast, we found no differences for these markers in tumors derived from 3-month-old fish (Supplementary Figure 3B-D). Nevertheless, GSEA analysis showed incipient profiles of reduction in cell proliferation and higher apoptosis even in early *tert-/-* tumors (Supplementary Figure 3A). Our results show that lack of telomerase affects both cell cycle progression and cell survival. This finding could explain the differences in tumor growth between *tert+/+* and *tert-/-* fish at a later stage of tumorigenesis. In addition, our expression profiles also show an increase in expression of genes related to melanocyte differentiation, suggesting that melanoma cells may exhibit a propensity to differentiate back to melanocytes. Consistently, melanocyte differentiation of melanoma cells was previously identified as one of the major expression signatures of regressing melanomas in the unique MeLiM minipig model^34^. Our data indicates that growth restriction of late *tert-/-* melanoma encompasses a tumor-autonomous effect by reducing cell proliferation, increasing apoptosis and melanocyte differentiation.

**Figure 4:**
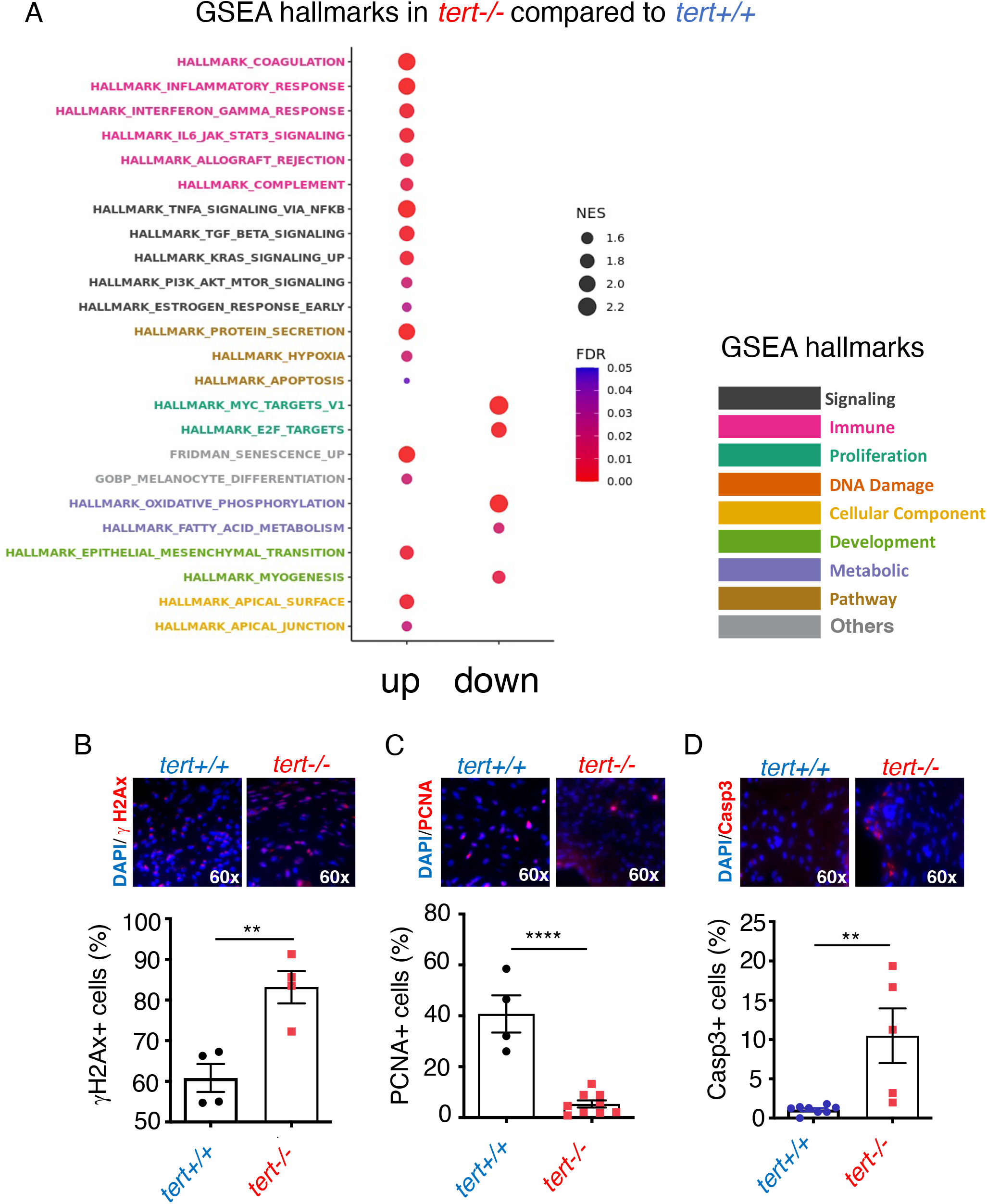
Lack of telomerase leads to low proliferation, increased apoptosis, inflammation and melanocyte differentiation. A-Gene Set Enrichment Analysis (GSEA) hallmarks of late stage melanoma of *tert-/-* fish compared to *tert+/+* highlighting inflammatory pathways, senescence and melanocyte differentiation. (B-D) Representative immunofluorescence images and respective quantifications of *tert+/+* and *tert-/-* tumors. B-DNA damage by γH2AX, C-proliferation by PCNA, D-apoptosis by cleaved-Caspase 3 immunofluorescence of 9-month-old fish tumors (unpaired *t*-test). Error bars represents ±S.E.M.; each dot represents an individual tumor; N≥4; * *p*≤0,05; ** *p*≤0,01; \*\*\*\**p*≤0,0001.

### Non-tumor autonomous causes for *tert-/-* tumor growth decline

GSEA analysis also revealed that up-regulation of the immune system was a hallmark of late *tert-/-* melanomas (Figure 4A). Interestingly, one of the inflammation GSEA pathways up-regulated in late tert-/- was allograft rejection (Figure 4A and 5A). Gene expression profiles consistent with immune system alterations were already present in early *tert-/-* tumors, despite ranking amongst the fourth-most significant cellular networks affected at that timepoint (Supplementary Figure 3A). In order to test whether *tert-/-* tumors were more immunogenic, we compared levels of immune infiltrates in melanoma of early and late tumors. We performed immunofluorescence staining for immune cells and this analysis revealed a robust tumor infiltration of the late stage *tert-/-* tumors (L-Plastin, *p*= 0,0310, Figure 5B), although some infiltration could be detected in early *tert-/-* tumors (*p*= 0,0493, Supplementary Figure 4A). Our results show that *tert-/-* melanoma is more immunogenic than tert+/+ tumors. Our data also suggests that an immune response may play a role in restricting the later stages of growth of *tert-/-* melanomas. In agreement, regressing melanoma of MeLiM minipigs was also shown to be accompanied by a robust immune response^34^.

**Figure 5:**
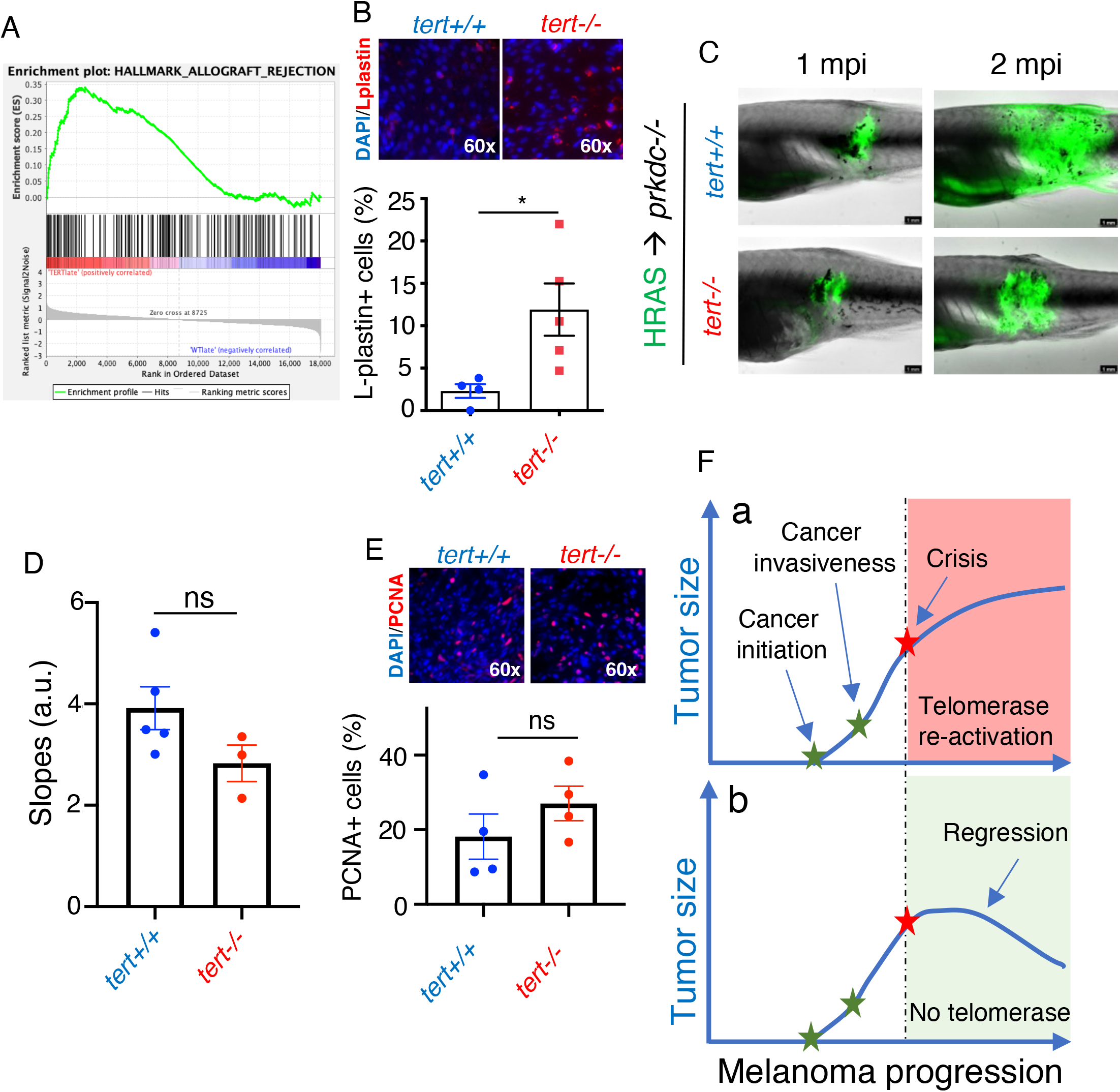
The immune system restrains growth of telomerase deficient melanoma. Telomerase deficient tumors are strongly immunogenic. A-Allograft rejection enrichment plot. B-Immunofluorescence images and quantifications of immune cell infiltrates (L-plastin) of late-stage melanoma. Late stage *tert-/-* melanoma is able to proliferate and grow in immunocompromised fish. C-Images of immunocompromised zebrafish (*prkdc-/-*) engrafted with primary melanoma cells derived from 9-month-old zebrafish at 1 month post-injection (mpi) and 2 mpi. D-Progression slope of tumor area calculated by a linear regression of two time-points (1mpi and 2mpi). E-Cell proliferation of tumor cells quantified using PCNA immunofluorescence (n≥ 4; unpaired *t*-test). F-We propose that melanoma can initiate and progress without telomerase activation. However, once telomeres become critically short, cancer cells must activate telomerase to sustain proliferation at later stages (a). If they fail to do so, tumor growth stagnates leading to tumor regression. Error bars represent ±S.E.M.; each dot represents an individual tumor; *** *p*≤0,001; ns: not significant.

To test the idea that the immune system could restrain the growth of *tert-/-* melanomas, we transplanted melanoma cells derived from 9 month-old tumors into immune-compromised *prkdc-/-* fish. These fish are deficient in DNApk and lack mature B an T cells^35^. We assessed tumor growth between 1 and 2 months after transplantation. Consistent with previous data, *tert+/+* tumors increased in size over this period (Figure 5C). Strikingly, despite a more restrained growth, *tert-/-* tumors also increased size when transplanted into immune-compromised fish (*p*=0,1312, Figure 5C-D). To show that the increase in tumor size was due to cell division, rather than simple cell expansion, we measured cell proliferation by immunofluorescence. Indeed, the rate of cell proliferation was similar between *tert+/+* and *tert-/-* tumors (*p*=0,2872, Figure 5E), denoting that *tert-/-* tumor cells were capable of sustained cell proliferation. Unfortunately, due to high mortality, we were unable to observe tumor growth beyond 2 months post-injection. Finally, to test whether the transplanted *tert-/-* tumors were not a selection of cells that activated a TMM, we measured C-circles to assess ALT activation. Similar to the original tumors in *tert-/-* fish, we did not detect any signs of ALT in the transplanted tumors (*p*>0,9999 compared to HeLa, Supplementary Figure 5A, B). Altogether, our results indicate that, even though *tert-/-* tumors stagnate and even regress in *tert-/-* fish, when transplanted to an immune-compromised environment, they can proliferate and grow in size. Thus, not only *tert-/-* tumors are more immunogenic but also they are restrained from growth by the organism adaptive immune system, similar to what has been observed in humans through CD8-positive cytotoxic T lymphocytes^34,36^.

## Discussion

The identification of activating TPMs in high frequency for several human cancers, including cutaneous melanoma^20,21,24^, highlights the role of telomere protection and TMMs in tumorigenesis. Previous studies provided evidence that TPMs may occur as an early step in carcinogenesis. This event would likely confer an advantageous evolutionary step rendering cell immortality to early cancer precursors. However, the absolute requirement of a TMM for early tumorigenesis is still unclear.

In our current study, absence of telomerase in two established zebrafish melanoma models (HRAS and BRAF) results in the same tumor incidence as WT controls. Additionally, early *tert-/-* tumors display similar invasiveness levels and tumor size as *tert+/+*. Thus, contrary to our initial expectation, telomerase is not required for melanoma initiation and early progression in zebrafish. These results are also surprising taking into consideration that late generation telomerase deficient mice (G5 terc-/-) are resistant to skin tumorigenesis^11^. A potential difference between the these models may reside in the fact that G5 terc-/- mice have considerably shorter telomeres from birth^11^. Thus, G5 terc-/- pre-cancer cells would attempt to proliferate with short telomeres capable of restricting cell proliferation. In contrast, *tert-/-* zebrafish possesses normal telomere length during development and differences are only visible in adult fish, as they result from *tert+/-* incrosses. Indeed, we fail to detect telomere shorting in early melanoma of either *tert+/+* or *tert-/-* fish. This may occur because melanoma, at this stage, have not yet undergone enough cell divisions to show an appreciable reduction of average telomere length when compared to skin controls. Alternatively, melanoma initiation in the mitfa:HRAS model may not have a single origin leading to multiple clones with different telomere length.

Similar to early *tert-/-* melanoma, we neither observe telomerase activity nor appreciable telomere shortening in *tert+/+* tumors. This finding is consistent with the similar tumor incidence observed in *tert+/+* or *tert-/-* fish, arguing that critical telomere shortening fails to restrain tumorigenesis at early stages of melanoma formation. This result differs from previous observations that link TPMs to an early event in tumorigenesis^30^. However, Chiba *et al*. demonstrated that, despite carrying TPMs, human melanoma cells still shorten telomere length with similar kinetics than other primary cells and that telomerase upregulation only occurs after many cell divisions concomitant with accumulation of critically short telomeres^30^. Thus, even though TPMs may occur early in tumorigenesis, telomere length in pre-cancer cells may be sufficient to sustain early cell divisions so that activation of telomerase and telomere elongation constitute a later event required to sustain continuous tumor growth. Consistent with this notion, Viceconte *et al*. identified a set of aggressive human melanoma metastasis that did not present either telomerase activity or ALT as TMMs^15^. Our work appears to replicate these findings of human melanoma and, thus, constitute a timely animal model to study cancers that fail to activate a TMM.

In contrast to early melanoma, we observed that telomerase becomes active at later stages, while its absence restricts tumor growth leading to tumor regression. These results reaffirm the requirement of a TMM in tumorigenesis, albeit at an unsuspected later stage of cancer progression. Nevertheless, melanoma results in zebrafish mortality, even in absence of TMMs. Ackermann *et al*. reported that half of studied neuroblastomas, a pediatric tumor of the sympathetic nervous system, also did not possess a TMM^37^. Neuroblastomas lacking TMMs were also associated with better prognosis characterized by non-progression and spontaneous regression.

Cases of human melanoma regression also occur at often unsuspected rates, accounting for 10 to 35% of the totality of melanomas^38^. However, the mechanism of human melanoma regression is currently unknown. We propose that telomere shortening may account for melanoma regression by imposing a two-fold barrier to tumor growth: 1-A “classical” tumor-autonomous mechanism whereby short telomeres restrain cell division while inducing apoptosis and melanocyte differentiation. Consistently, melanoma regression in other models is also accompanied by cell cycle arrest, apoptosis and melanocyte differentiation^34,39^, even though telomere shortening was not assessed as a potential cause. 2-A second unsuspected barrier consisting of a non-tumor autonomous mechanism imposed by the immune system. *tert-/-* tumors are more immunogenic possessing higher numbers of immune cell infiltrates. Consistently, GSEA analysis of *tert-/-* tumors highlighted upregulation of immune response pathways, including terms such as allograft rejection. Conclusively, *tert-/-* tumors were able to proliferate and grow once transplanted into immunocompromised fish, lacking B and T cells. This second barrier was previously observed in regressing melanoma of MeLiM minipigs and humans, in particular through CD8-positive cytotoxic T lymphocytes^34,36^. In agreement with our data, human cancers harboring mutations in TERT were previously highlighted as immunogenic, leading to a better survival upon immunotherapy^18,19^. In agreement, melanoma susceptible to immune checkpoint therapy had less expression of genes related to telomere maintenance^17^. Conversely, TPMs leading to telomerase expression were associated with resistance to immunotherapy in metastatic renal cell carcinoma, being proposed as a negative predictor of outcome for immunotherapy^40^. Altogether, these studies suggest that telomerase expression protects tumors from being a target of the immune system.

In summary, our data shows that telomerase is not required for initiation and early progression of melanoma in zebrafish (Figure 5E-a). However, once telomeres reach a critical threshold of erosion, cells re-activate telomerase to sustain cell proliferation and cancer growth. If they fail to do so, telomeres continue to shorten (Figure 5E-b), and tumors stagnate growth and even regress. This is caused by two mechanisms: a tumor-autonomous mechanism (cell cycle arrest, apoptosis and melanocyte differentiation) and a non-tumor-autonomous mechanism (immune response). Our results illuminate an unappreciated contribution of the immune system in tumor formation in telomerase-negative tumors. We found that, despite the apparent lack of dependency on a TMM in tumor initiation and early progression, later stages of melanoma growth are influenced by immune function. It remains to be tested if immune activation in human tumors with critically eroded telomeres may enhance the ability to impair tumor growth.

## Methods

### Ethics statement

All experiments involving animals have been approved in France by the Animal Care Committee of the IRCAN, the regional (CIEPAL Cote d’Azur #784) and national (French Ministry of Research #33396-2021112517136391) authorities and in Portugal by the Ethical Committee of the Instituto Gulbenkian de Ciência and approved by the competent Portuguese authority (Direcção Geral de Alimentação e Veterinária; approval number: 0421/000/000/2015).

### Zebrafish maintenance

Zebrafish were maintained in accordance with Institutional and National animal care protocols. The telomerase mutant zebrafish *tert* ^+/hu3430 26^(referred in this work as *tert+/-*) were crossed either with *Tg(mitfa:Hsa.HRAS_G12V,mitfa:GFP)* ^27^or *Tg(mitfa:BRAFV600E); tp53^+/M214K^* ^28^ to generate the stock melanoma lines containing the *tert* mutation. All *tert+/-* lines are maintained as outcross to *tert+/+* to avoid the effects of haploinsufficiency in the progeny^41^ The experimental fish were obtained by crossing *tert+/-* fish with either *Tg(mitfa:Hsa.HRAS_G12V,mitfa:GFP); tert+/-* or *Tg(mitfa:BRAFV600E); tp53^+/M214K^; tert+/-* to ensure that HRAS+ and BRAF+ fish possessed a single copy of the oncogene. Finally, to generate immunocompromised zebrafish, we incrossed the *prkdc*^+/D3612fs 35^ line to generate *prkdc^-/-^* homozygous fish.

### Tumor size quantification

Fish were anesthetized with 160μg/L of tricaine methane sulfonate solution (MS222), placed on a 1% agarose-coated petri dish and immediately imaged using a fluorescence stereomicroscope (Leica M205FA). After the procedure, fish were placed back in tanks containing system water and monitored for recovery until they regained a normal swimming pattern. Images were analyzed by the software ImageJ (version 2.9.0). We outlined the tumor and quantified its area by considering both the GFP signal and visible outgrowth on the skin. To determine tumor growth, we generated a linear regression of the area at different timepoints and obtained a slope to represent progression.

### Fixation for histology and immunofluorescence

The animals were euthanized with 1g/L of MS222 and then fixed in 10% neutral buffered formalin for 72 hours at room temperature. The fish were then decalcified in 0.5M EDTA for 48 hours and the whole fish were embedded in paraffin. Transversal sections of 5μm were obtained, and the slides were stained with hematoxylin and eosin for histopathological analysis. Additionally, slides were used for immunofluorescence analysis.

### Immunofluorescence of paraffin embedded zebrafish

Paraffin-embedded sections were deparaffinized with HistoChoice® clearing agent (H2779, Sigma-Aldrich) and rehydrated using a series of alcohols (100%, 80%, 60%). Slides were then placed in citrate buffer (10 mM, pH 6) and microwaved at 550W for 20 minutes. After cooling, slides were washed in PBS followed by incubation in blocking buffer (2% Normal Goat Serum, 1% DMSO, 0,1% tween in PBS) for 1 hour at room temperature. Subsequently, slides were incubated overnight at 4°C with primary antibody diluted in blocking buffer (these were: 1:200 Living Colors Ab, #632380, Takara; 1:100 L-plastin, #GTX124420, GeneTex; 1:50 γH2AX phosphoSer139; #GTX127342; GeneTex; 1:50 Cleaved Caspase-3, #ab13847, Abcam; 1:50 PCNA, #sc7907, Santa Cruz). The following day, slides were washed 3 times with PBS and incubated with the secondary antibody (1:500) for 2 hours at room temperature. This was followed by 3 washes with PBS and incubated with DAPI (1:2000 in dH2O) for 10 minutes. Finally, slides were washed with PBS for 10 minutes at room temperature and mounted with Fluoroshield™ (#F6182, Sigma). Stainings were imaged on Delta Vision Elite (GE Healthcare). Whole tumor area was identified by the GFP staining. Quantification of the different antibodies was performed by quantifying the percentage of positive cells within the tumor area for a minimum of 200 cells per slide.

### Sample collection

The fish with tumors were euthanized using 1g/L of MS222, and the tumors and/or skin were dissected and rinsed with ice-cold PBS. Samples intended for molecular analysis were either used immediately or snap-frozen in liquid nitrogen and stored at –80°C. Tumor samples designated for allografts were utilized without further processing.

### Telomere restriction fragment (TRF) analysis by Southern blot

Tumor and skin samples were lysed overnight at 50°C in lysis buffer (#K0512, Fermentas) with 1 mg/ml Proteinase K and RNase A (1:100) and genomic DNA was extracted using a standard phenol-chloroform protocol. Genomic DNA was then digested with RSAI and HINFI enzymes for 12 hours at 37°C. After digestion, samples were loaded on a 0.6% agarose gel in 0.5% TBE buffer. The gel was run at the constant voltage of 110V for at least 17 hours at 4°C. Gels were then fixed with 0.25N HCl for 15 minutes and processed for Southern blotting using a 1.6 kb telomere probe, (TTAGGG)n, labelled with [α-32P]-dCTP.

### C-circle assay

C-circle assay was carried on as described by Henson *et al*. ^42^. Briefly, 30ng of genomic DNA was used for each reaction containing 7.5 U Φ29 (#M0269L, BioLabs), 1x Φ29 DNA polymerase buffer, 0.2 mg/ml BSA, 0.1% (v/v) Tween 20, 1 mM each dATP, dTTP, dGTP, 4 μM dithiothreitol (DTT). The rolling circle amplification reaction proceeded as follow: 8 hours at 30°C and 20 minutes at 65°C using a thermocycle machine. For each sample, a reaction without the Φ29 DNA polymerase was used as a control (-Φ29). Using a 96-well Bio-Dot^TM^ Apparatus (Bio-Rad), reaction products (1:8 dilution in 2x SCC) were dot-plotted onto a 2x SCC pre-soaked positive nylon membrane. The membrane was then UV-crosslinked and hybridized with a 1.6 kb telomere probe, (TTAGGG)n, labelled with [α-32P]-dCTP.

### *TRAP* assay

Tumor samples were homogenized in 0.5% CHAPS followed by incubation on ice for 30 minutes. Samples were then centrifuged (16000xg for 20 min at 4°C) and the supernatant was collected. Protein concentration was measured using a Bradford assay according to manufacturer’s instructions. 0.5 μg of whole protein was added to the TRAP master mix, which contained 1x HotStart Taq Polymerase (#203205, Qiagen), 1x TRAP buffer (20mM Tris-HCL pH 8.3, 1.5mM MgCl2, 63mM KCl, 0.05% (v/v) Tween 20 and 1mM EGTA), 25μM dNTPs, 0,8 ng/μl BSA and the following primers 100ng/μl ACX: GCGCGGCTTACCCTTACCCTTACCCTAACC, 100ng/μl NT: ATCGCTTCTCGGCCTTTT, 0.01 attomol/μl TSNT: AATCCGTCGAGCAGAGTTAAAAGGCCGAGAAGCGAT, 2ng/μl TS: AATCCGTCGAGCAGAGTT. As negative control, samples were treated with 12.25mg/ml Rnase for 20 minutes at 37 °C. Reactions were then kept in the dark for 30 minutes at room temperature, followed by 15 min at 95°C, 30 cycles of 30s at 95°C, 30s 52°C and 45s at 72°C and a final step of 10 min at 72 °C. TRAP products were run on a 10% polyacrylamide gel with 1x TBE.

### RNA extraction/ RNA sequencing

RNA extraction was performed using a TRIzol-chloroform protocol according to manufacturer’s specifications. RNA quality was assessed by BioAnalyzer (Agilent 2100). RNA-seq library preparations and sequencing were outsourced to Beijing Genomics Institute (BGI, Hongkong). Briefly, pair ended 150 bases reads were sequenced on DNBseq platform and 100M clean reads per sample was generated. The RNA-seq data was analyzed by an internal pipeline for transcript comparison. Before running, genes were mapped to zebrafish orthologs using Ensembl’s Biomart database. Using MsigDB’s Hallmark gene set^43,44^, the gene set enrichment analysis (GSEA) was run on the Broad Institute’s GSEA Software. One thousand permutations were run with a gene set permutation type. Significant enrichment results (p≤0.05) were considered to be GSEA hallmarks. Borderline significance (to show directionality) was set at nominal p-values below 0.05 and false discovery rate below 0.25.

### Tumor allografts

Tumor samples were dissociated with TryPLe (ThermoFisher Scientific) for 20 minutes at 30°C and filtered using a 70μm cell strainer (#22363538, Fisher Scientific). Cells were then stained with Trypan Blue and counted with a hemocytometer and diluted in PBS to a final concentration of 100 000 cells/μl. Immunocompromised zebrafish (*prkdc-/-*) were anesthetized with 160μg/L MS222 and injected subcutaneously with ca. 500 000 cells on the left flank below the dorsal fin. After the procedure, fish were placed back into tanks containing system water and their recovery was monitored until they regained normal swimming pattern. Injected fish were imaged using a fluorescence stereomicroscope (Leica M205FA) at 1- and 2-months post injection.

### Telomere Dysfunction-Induced Focus (*TIF*)

The tumors were excised and dissected to small pieces. They were then transferred to a solution of 1.1% sodium citrate and left at room temperature for 8 minutes, followed by an additional 8 minutes on ice. After this, the samples were transferred to a 3:1 solution of methanol and acetic acid. The solution was replaced after 20 minutes, and the samples were kept overnight at –20°C. The following day, the samples were transferred to a 50% acetic acid solution, the tissue was dissociated with a blade, and the cells were filtered through a 70μm cell strainer (#22363538, Fisher Scientific). The cells were then deposited onto a slide using a cytospin centrifuge and allowed to air-dry. The cells were permeabilized with 0.5% Triton-X-100 for 8 minutes and dehydrated with a series of ethanol. Next, hybridization of the telomere PNA probe (#F1002, Panagen) was performed for 2 hours at room temperature after denaturation in a solution of 10mM Tris-HCl pH 7.2, 70% formamide, and 1% blocking solution (#11096176001, Roche) for 3 minutes at 85°C. The cells were then washed for 30 minutes with 10mM Tris-HCl pH 7.2, 70% formamide, followed by a second wash with 50mM Tris-HCl pH 7.5, 150 mM NaCl for 15 minutes. The cells were then incubated with a blocking buffer (1% Triton-X-100, 1% BSA, 5% normal goat serum) for 1 hour, followed by primary antibody incubation (1:50 γH2AX phospho Ser139; #GTX127342; GeneTex) overnight at 4°C. The cells were then washed with PBS, followed by secondary antibody incubation for 2 hours at room temperature and 10 minutes of DAPI incubation (1:2000 in dH2O). Finally, the slides were mounted with FluoroshieldTM (#F6182, Sigma), and the staining was imaged on a Delta Vision Elite (GE Healthcare). The number of γH2AX foci colocalizing to telomeres was quantified per nucleus. A minimum of 100 nuclei were assessed per group.

### Statistical analysis

Statistical analysis and graphs were done in GraphPad Prism 8.4.0 software. The specific statistical test used for each graph is described in the corresponding figure legend. A *p*≤0.05 was considered significant throughout of the study.

## Data availability

All data generated or analysed during this study are included in this published article and its Extended information files. The raw RNA-seq reads generated in this study have been deposited to the SRA database under the accession number XXX.

## Acknowledgments

We are grateful to Lea Harrington (University of Toronto, Canada) and Elizabeth Patton (University of Edinburgh, UK) for critically reading our manuscript. We also thank Richard White for kindly providing the ZMEL cells. This work was supported by the Portuguese Fundação para a Ciência e a Tecnologia (FCT, PTDC/BIM-ONC/3402/2014), Université Co□te d’Azur - Académie 4 (Installation Grant: Action 2 - 2019), Fondation ARC pour la Recherche sur le Cancer (PJA20161205137), Institut National du Cancer (INCa, PLBIO21-228) and the Howard Hughes Medical Institute International Early Career Scientist grant awarded to M.G.F. B.L.-B. was supported by a postdoctoral fellowship awarded by the Fondation pour la Recherche Médicale (FRM, SPF201809007006). We thank the Instituto Gulbenkian de Ciência (IGC) histology unit, the IGC imaging unit and the IGC Fish Facility for excellent animal care, for assistance with experimental planning, sample and data collection. IGC Fish Facility is financed by Congento LISBOA-01-0145-FEDER-022170, co-financed by FCT (Portugal) and Lisboa2020, under the PORTUGAL2020 agreement (European Regional Development Fund). The work was also performed using the PEMAV fish facility, Imaging core facility (PICMI) and the Genomics facilities at the IRCAN supported by FEDER, Région Provence Alpes-Côte d’Azur, Conseil Départemental 06, ITMO Cancer Aviesan (plan cancer), Cancéropole Provence Alpes-Côte d’Azur, Gis Ibisa, CNRS and Inserm. The funders had no role in study design, data collection and analysis, decision to publish or preparation of the manuscript.

## Author contributions

B.L.-B. performed the experiments and carried out data analyses; J.N. and T.F. performed the BRAF experiments; T.F. and M.E.M. performed and analyzed TRFs; S.T. quantified IFs; J-X.Y., D.K. and M.B. performed transcriptomics analyses; T.C. performed histopathology analyses; B.L.-B. and M.G.F. conceived the study, designed the experiments, and wrote the manuscript. M.G.F. supervised the work.

## Declaration of competing interests

The authors declare no competing interests.

---

**Supplementary Figure 1: Telomerase does not contribute to melanoma initiation and early invasiveness.** (A-B) Lack of telomerase does not restrict melanoma incidence of BRAFV600E zebrafish. A-Percentage of tumor-free fish compared between *tert+/+* and *tert-/-* fish (log-rank test). B-Representative images of hematoxylin & eosin staining of different stages of melanogenesis. (C-F) Incidence of microscopic tumors is unaffected by the absence of telomerase. C-Cumulative incidence of microtumors of mitfa:HRAS fish (n=6/ timepoint; χ^2^ test). D-Representative images of Hematoxylin & Eosin staining of *tert+/+* and *tert-/-* microtumors. Black arrows identify microtumors. E-Cumulative percentage of tumors with invasive melanoma (n=6/timepoint; χ^2^ test). F-Representative images of hematoxylin & eosin staining of *tert+/+* and *tert-/-* invasive melanoma. Black arrows show melanoma invading muscles tissue. ns: not significant.

**Supplementary Figure 2: Lack of telomerase leads to melanoma regression but does not affect mortality.** Absence of telomerase leads melanoma regression. A-Representative images of melanoma regression in *tert-/-* fish. Melanoma mortality is not affected by the availability of telomerase. B-Kaplan-Meier survival of *tert+/+* and *tert-/-* fish melanoma either carrying the HRAS oncogene (full lines) and their control cousins (dashed lines). C-Scheme of crosses to generate the experimental fish and its controls.

**Supplementary figure 3: Telomerase deficiency impacts gene expression of early stage melanoma.** A-Gene Set Enrichment Analysis (GSEA) hallmarks of late stage melanoma of *tert-/-* fish compared to *tert+/+*. (B-D) Representative immunofluorescence images and respective quantifications of *tert+/+* and *tert-/-* tumors. B-DNA damage by γH2AX, C-proliferation by PCNA, D-apoptosis by cleaved-Caspase 3 immunofluorescence of 3-month-old fish tumors (unpaired *t*-test).Error bars represent ±S.E.M.; each dot represents an individual tumor; N≥4; * *p*≤0,05; ns: not significant.

**Supplementary figure 4: Early-stage tert-/- melanoma displays infiltration of immune cells. A-** Immunofluorescence images and quantifications of immune cell infiltrates (L-plastin) of *tert+/+* and *tert-/-* tumors derived from 3-months old fish (unpaired *t*-test). Error bars represent ±S.E.M.; each dot represents an individual tumor; N≥4; * *p*≤0,05.

**Supplementary figure 5: Late melanoma allotransplants do not engage in ALT.** Even though *tert-/-* melanoma cells can proliferate in an immune compromised environment, they do not engage in ALT. A-C-circle analysis of 2mpi transplanted tumors. Extracts from HeLa cells and U2OS were used as negative and positive controls, respectively. B-Quantification of C-circle assay (n=5). Error bars represent ±S.E.M.; each dot represents an individual tumor; \*\*\**p*≤0,001; ns: not significant.

## Notes

### Competing Interest Statement

The authors have declared no competing interest.

