## Supplemental Figures for "Absence of Telomerase Leads to Immune Response and Tumor Regression in Zebrafish Melanoma"

Figure S1: Telomerase does not contribute to melanoma initiation and early invasiveness.

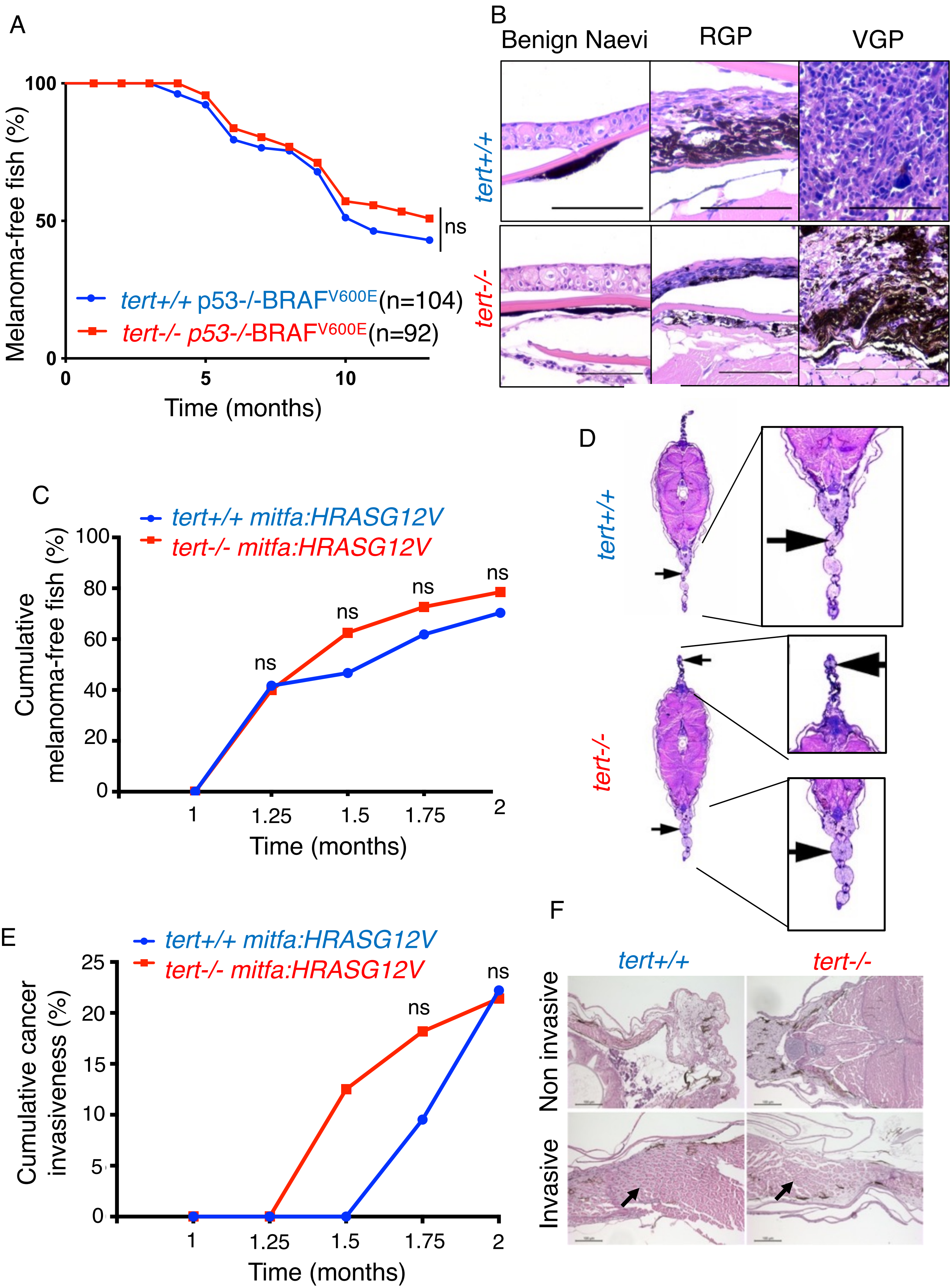

Figure S2: Lack of telomerase leads to melanoma regression but does not affect mortality.

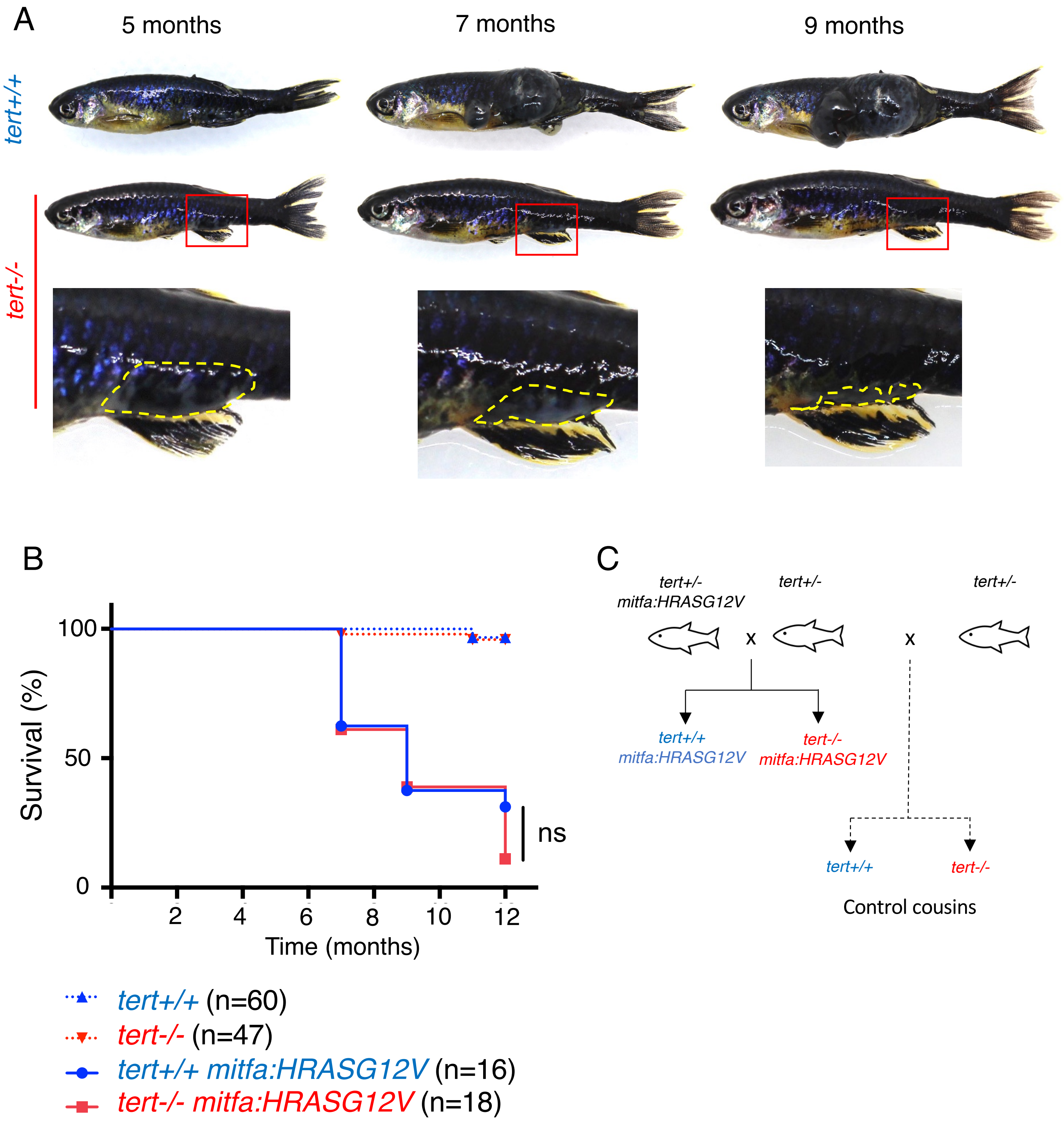

Figure S3: Telomerase deficiency impacts gene expression of early stage melanoma.

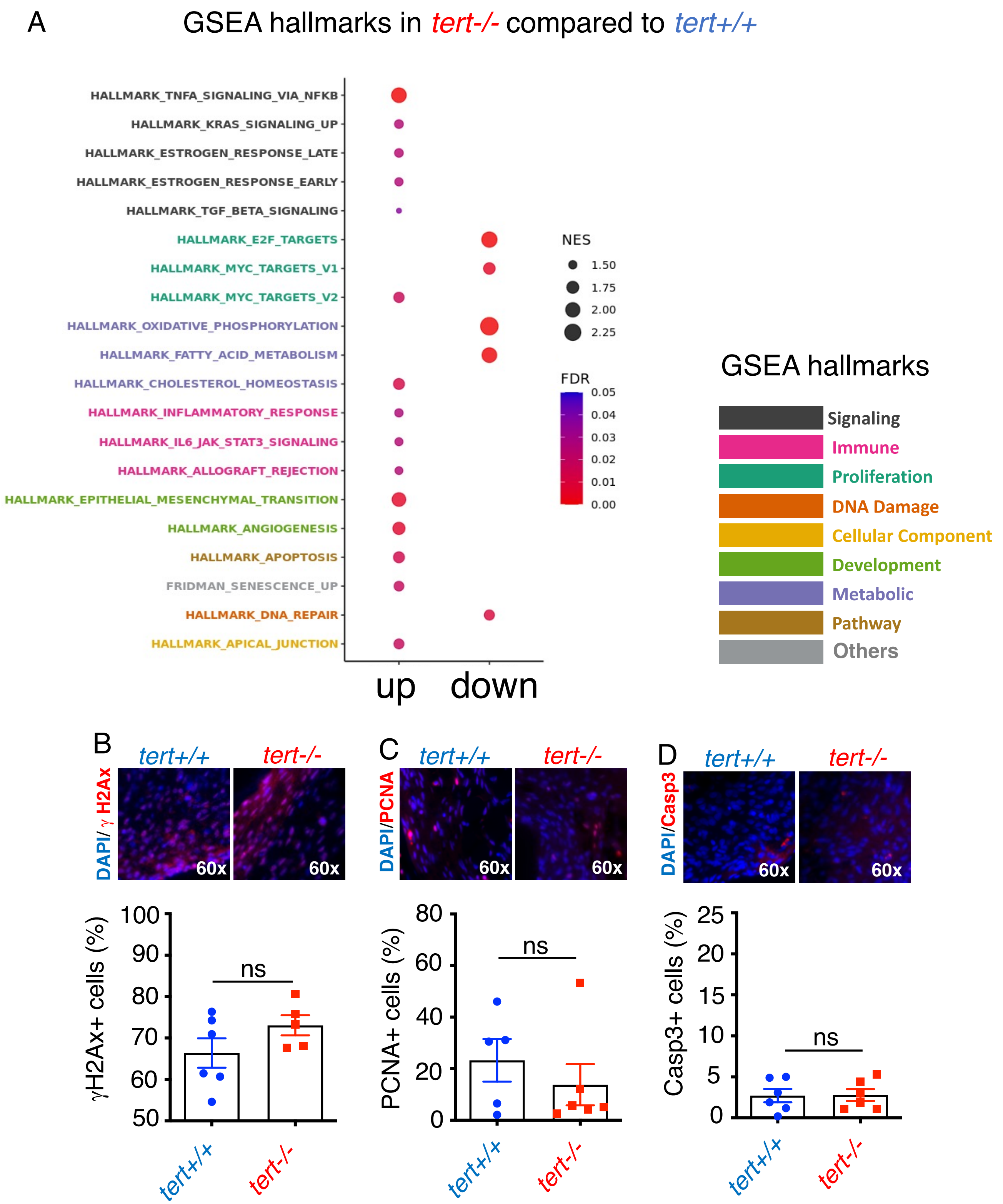

Figure S4: Early-stage tert-/- melanoma displays infiltration of immune cells.

A

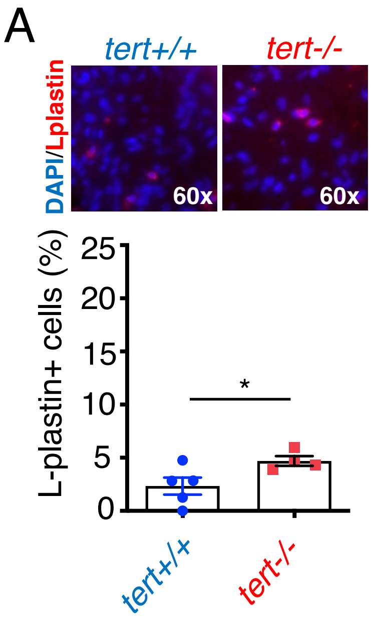

Figure S5: Late melanoma allotransplants do not engage in ALT.

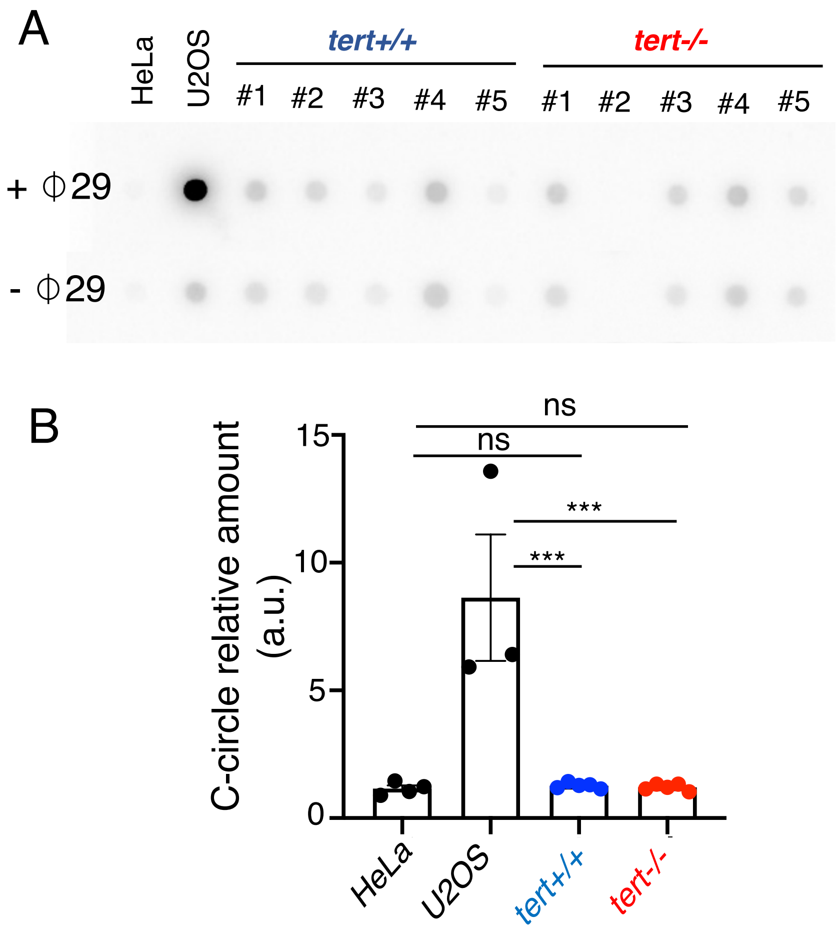
